# Triazolopyrimidines target aerobic respiration in *Mycobacterium tuberculosis*

**DOI:** 10.1101/2021.10.18.464924

**Authors:** Catherine Shelton, Matthew McNeil, Lindsay Flint, Dara Russell, Bryan Berube, Aaron Korkegian, Yulia Ovechkina, Tanya Parish

## Abstract

We previously identified a series of triazolopyrimidines with anti-tubercular activity. We determined that *Mycobacterium tuberculosis* strains with mutations in a component of the cytochrome *bc_1_* system (QcrB) were resistant to the series. A cytochrome *bd* oxidase deletion strain was also more sensitive to this series. We isolated resistant mutants, all of which had mutations in Rv1339. Compounds were active against intracellular bacteria but did not inhibit mitochondrial respiration in human HepG2 cells. These data are consistent with triazolopyrimidines acting via inhibition of *M. tuberculosis* QcrB.

---

We previously identified a series of triazolopyrimidines with anti-tubercular activity (1); compounds were bacteriostatic for replicating *Mycobacterium tuberculosis*, but bactericidal for non-replicating bacteria. We explored the structure-activity relationship and determined druglike properties. We wanted to determine the target and/or mechanism of action of the TZP series. Since previous work in our group and others had identified several common targets, we tested a set of analogs for activity against strains carrying mutations in promiscuous targets (Figure 1) (2).

**Figure 1.**
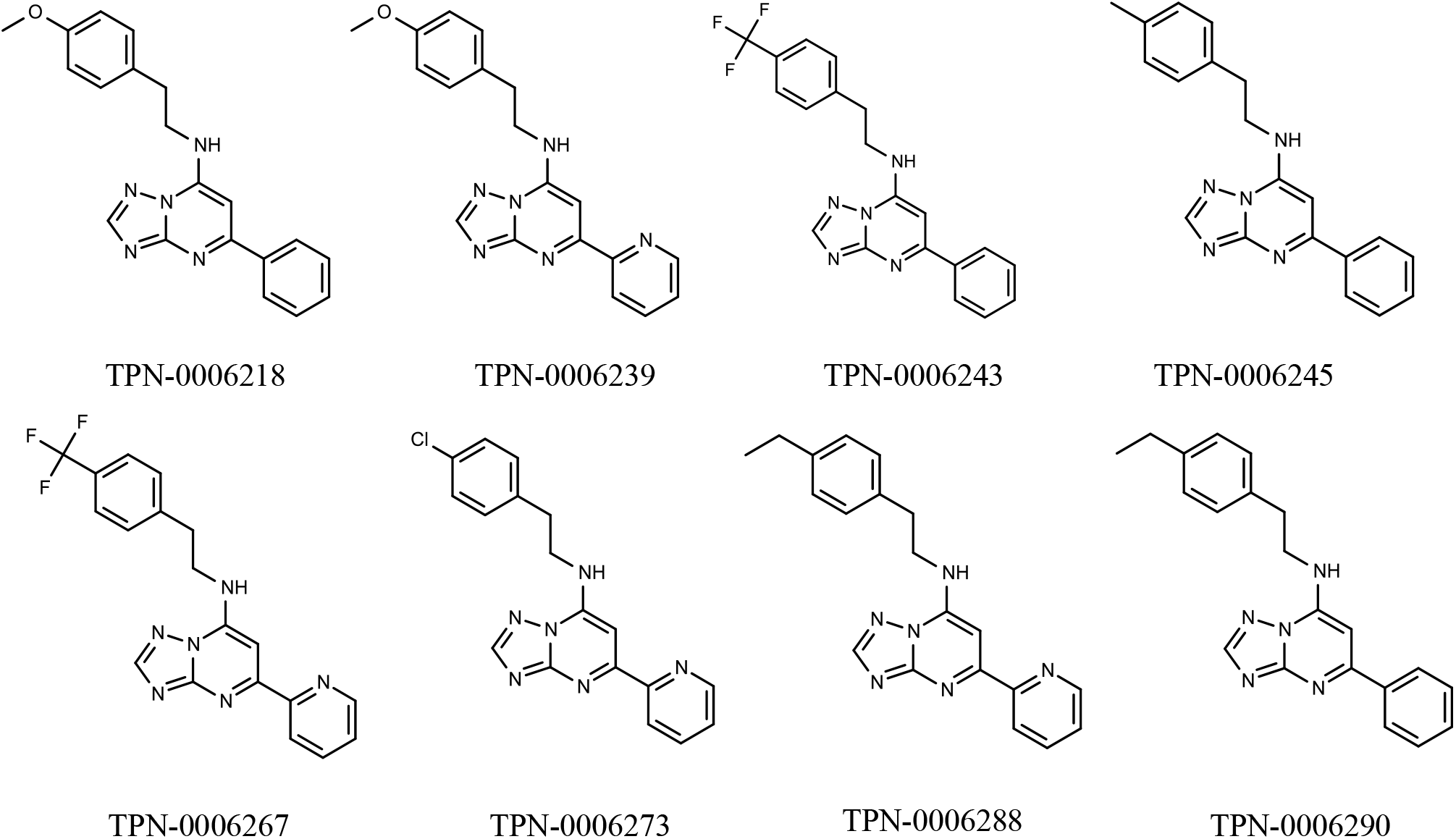
Structures of molecules.

We selected DprE1, MmpL3 and QcrB as the most common targets (3–6). We determined activity against strains carrying either DprE1_C387S_, MmpL3_F255L_ or QcrB_A396T_ mutations in the parental strain *M. tuberculosis* H37Rv-LP (ATCC 25618) (Table 1) (7). MICs were determined as described after 5 days growth in Middlebrook 7H9 medium plus 10% v/v OADC supplement and 0.05% w/v Tween 80 (8). We saw a small shift in MICs in the QcrB_A396T_ mutant strain of ~2 to 4-fold increase in resistance. No significant changes were seen in the DprE1_C387S_ or MmpL3_F255L_ mutant strains.

**Table 1.**
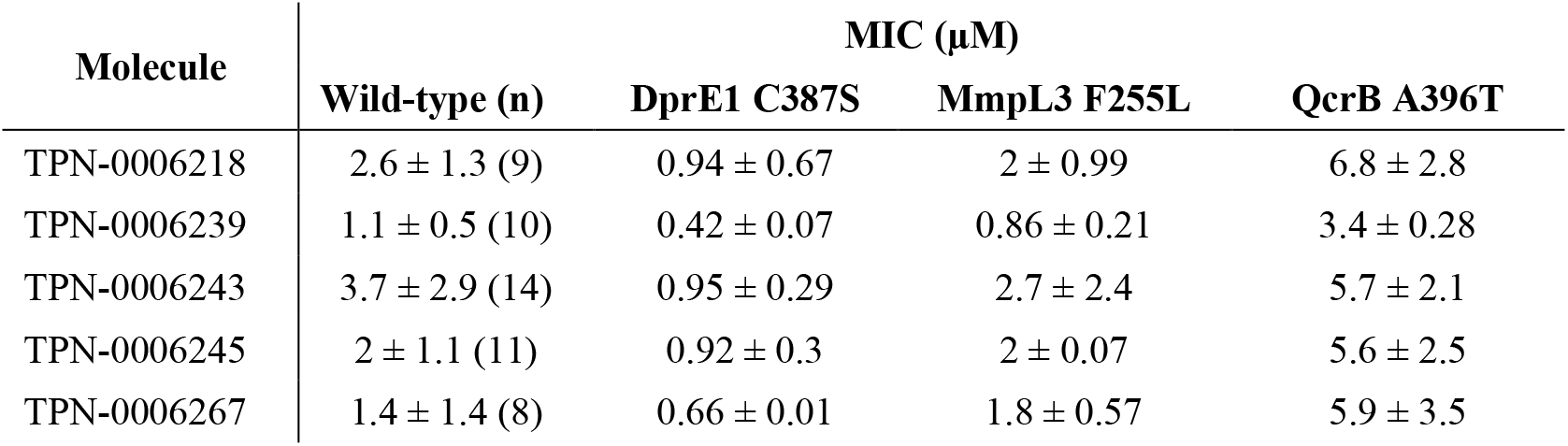
Activity against strains of *M. tuberculosis*. MICs were determined after 5 days in duplicate (except for wild-type where n = number of replicates). The genotype of the strain is noted. Wild-type is *M. tuberculosis* H37Rv-LP (ATCC 25618).

In order to confirm that QcrB mutation did lead to resistance and is the likely target, we tested compounds against two additional strains carrying QcrB mutations (T313I and M342T) (5,9), QcrB_T313I_ is the most commonly mutation which confers resistance to inhibitors (Table 2). We confirmed high level resistance in both strains.

**Table 2.**
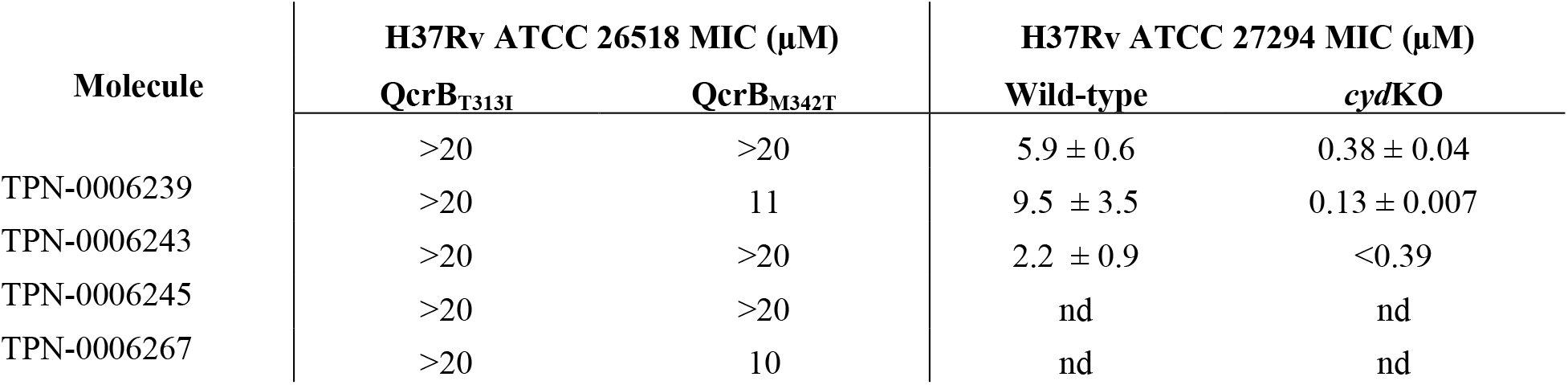
Activity against strains of *M. tuberculosis*. MICs were determined after 5 days. The genotype of the strain and parental strain is noted. nd – not determined.

QcrB is a component of the cytochrome *bc_1_* complex in the electron transport chain; *M. tuberculosis* strains in which the alternative cytochrome oxidase (cytochrome *bd*) is deleted are hypersusceptible to QcrB inhibitors (10,11). We also tested three compounds against *M. tuberculosis* H37Rv ATCC 272942 and the isogenic CydC deletion strain (11). As expected, *M. tuberculosis* H37Rv ATCC 27294 was more resistant to the compounds than H37Rv ATCC 25618, as has been noted with other QcrB inhibitors, although the mechanism behind this is unknown (10–12). Deletion of cytochrome *bd* activity resulted in higher sensitivity to the three compounds (Table 2). Taken together these data strongly support the hypothesis that the target of the series is QcrB.

We wanted to determine if there were additional targets or mechanism(s) of resistance, so we isolated and characterized resistant mutants to the series. We selected compounds from our original set with the lowest liquid MIC and determined MIC against *M. tuberculosis* H37Rv ATCC 25618 on solid medium (Table 3). We selected two compounds and plated ~10^8^ bacteria onto 5X and 10X solid MIC as described (4). We isolated colonies and confirmed resistant mutants by measuring the MIC on solid medium; we obtained nine resistant isolates for TPN-0006239 and five resistant isolates for TPN-0006267 (Table 3).

**Table 3.** Characterization of resistant isolates of *M. tuberculosis*. MICs were determined in 24-well agar plates after 3 weeks of incubation. Two genes (*qcrB* and *rv1339*) were sequenced in all strains.

| Strain | Compound | MIC<br>(μM) | Rv1339 | QcrB |
| --- | --- | --- | --- | --- |
| H37Rv-LP | TPN-0006239 | 1.6 | wt | wt |
| LP-0497553-RM1 | TPN-0006239 | 25 | P121L | wt |
| LP-0497553-RM2 | TPN-0006239 | 25 | P121L | wt |
| LP-0497553-RM4 | TPN-0006239 | 50 | P121L | wt |
| LP-0497553-RM5 | TPN-0006239 | 50 | P121L | wt |
| LP-0497553-RM10 | TPN-0006239 | 50 | S120P | wt |
| LP-0497553-RM11 | TPN-0006239 | 50 | P121L | wt |
| LP-0497553-RM14 | TPN-0006239 | 50 | wt | wt |
| LP-0497553-RM15 | TPN-0006239 | 50 | P121L | wt |
| LP-0497553-RM23 | TPN-0006239 | 50 | wt | wt |
| H37Rv-LP | TPN-0006267 | 1.6 | wt | wt |
| LP-0499227-RM1 | TPN-0006267 | > 100 | P121L | wt |
| LP-0499227-RM2 | TPN-0006267 | > 100 | P121L | wt |
| LP-0499227-RM3 | TPN-0006267 | > 100 | P121L | wt |
| LP-0499227-RM4 | TPN-0006267 | 25 | P121L | wt |
| LP-0499227-RM7 | TPN-0006267 | > 100 | P121L | wt |

We sequenced the entire QcrB gene in all fourteen, but none of them had mutations (Table 3). We had previously seen mutations in Rv1339 leading to resistance to other QcrB inhibitors (5,9), so we sequenced Rv1339. We found the same mutation in eleven strains (P121L); one strain had the mutation S120P (Table 3). Two strains had no mutations in Rv1339. We have previously linked Rv1339 mutations to resistance to other QcrB inhibitor series, namely the imidazopyridines and the phenoxyalkylimidazoles (5,9). Recent work in the related organism *Mycobacterium smegmatis* suggests that Rv1339 is an atypical class II cAMP phosphodiesterase that has been linked to antibiotic tolerance (13). In addition a P94L mutation in Rv1399 led to increased persistence in animal models and increased resistance to external stress in *Mycobacterium canetti*, proposed to be due to changes in cell wall permeability (14). It is possible that the mutations we obtained lead to decreased compound permeation leading to resistance. However, it is unusual that resistance is largely seen with QcrB inhibitors, not as a general phenomenon; an alternative explanation for resistance could be changes in the intracellular ATP pool due to decreased turnover of cAMP.

We had previously demonstrated that this series had bacteriostatic activity against replicating *M. tuberculosis*, but bactericidal activity against non-replicating bacteria (1). We have noted this biological activity profile for other QcrB inhibitors and thus it is consistent with it being an inhibitor of aerobic respiration (5,9,12). Since other QcrB inhibitors are active against intracellular bacteria, we tested the TZP series for activity against *M. tuberculosis* in human THP-1 macrophages. Macrophages were infected with *M. tuberculosis* expressing luciferase (15) at a multiplicity of infection of ~1 for 24h, washed to remove extracellular bacteria, and then exposed to compound for 72 h. Bacterial growth was measured by fluorescence. We tested five representative molecules, and all had potent activity with IC_50_ < 1 μM (Table 4).

**Table 4.** Activity against intracellular *M. tuberculosis*. IC5_0_ were measured after 72h in THP-1 cells infected at an MOI of 1 (n=2).

| Molecule | Intracellular<br>$IC_{50}$ |
| --- | --- |
| TPN-0006218 | $0.23 \pm 0.08$ |
| TPN-0006267 | $0.21 \pm 0.11$ |
| TPN-0006273 | $0.19 \pm 0.13$ |
| TPN-0006288 | $0.076 \pm 0.032$ |
| TPN-0006290 | $0.18 \pm 0.09$ |

Since we identified the target of the TZP series as aerobic respiration, we determined whether the series might also inhibit mitochondrial respiration. We determined cytotoxicity against HepG2 cells cultured in DMEM with galactose as the carbon source to force the cells into using mitochondrial respiration (16). HepG2 cells were exposed to compound for 72 h and viability measured using CellTiterGlo (Promega) (1). Of eight compounds, six showed some cytotoxicity (Table 5), although they still had a good selectivity index (activity was more potent against *M. tuberculosis*). We compared the IC_50_s under this condition to those generated when HepG2 cells were cultured in glucose when mitochondrial respiration is not active (1). There was less than two-fold difference in the cytotoxicity, confirming that molecules are not inhibiting eukaryotic respiration.

**Table 5.** Cytotoxicity against human HepG2 cells. HepG2 cells were cultured in medium containing either galactose or glucose as the carbon source. IC_50_ = the concentration required to reduce cell number by 50% was determined after 3 days exposure to compounds.

| Molecule | IC<sub>50</sub> (μM) |  |
| --- | --- | --- |
|  | Glucose | Galactose |
| TPN-0006218 | >100 | 65 |
| TPN-0006239 | >100 | 73 |
| TPN-0006243 | >100 | >100 |
| TPN-0006245 | 58 | 39 |
| TPN-0006267 | >100 | >100 |
| TPN-0006273 | 100 | 76 |
| TPN-0006288 | 44 | 23 |
| TPN-0006290 | 49 | 33 |

In conclusion, we have determined that the triazolopyrimidine series inhibits *M. tuberculosis* growth by targeting QcrB, a component of the electron transport chain. In addition, we have demonstrated that mutations in either the target QcrB, or the putative phosphodiesterase Rv1339 lead to resistance. Since QcrB is a clinically validated target, this is an attractive series to develop further.

## Acknowledgements

We thank Lena Anoshchenko, Junitta Guzman, David Roberts, Dean Thompson and James Vela for technical assistance.

